# Quantifying the subjective cost of self-control in humans

**DOI:** 10.1101/2020.10.15.341354

**Authors:** Candace M. Raio, Paul W. Glimcher

## Abstract

Since Odysseus committed to resisting the Sirens, mechanisms to limit self-control failure have been a central feature of human behavior. Psychologists have long argued that the use of self-control is an effortful process and, more recently, that its failure arises when the cognitive costs of self-control outweigh its perceived benefits. In a similar way, economists have argued that sophisticated choosers can adopt “pre-commitment strategies” that tie the hands of their future selves in order to reduce these costs. Yet, we still lack an empirical tool to quantify and demonstrate the cost of self-control. Here, we develop and validate a novel economic decision-making task to *quantify the subjective cost of self-control* by determining the monetary cost a person is willing to incur in order to eliminate the need for self-control. We find that humans will pay to avoid having to exert self-control in a way that scales with increasing levels of temptation and that these costs are modulated both by motivational incentives and stress exposure. Our psychophysical approach allows us to index moment-to-moment self-control costs at the within-subject level, validating important theoretical work across multiple disciplines and opening new avenues of self-control research in healthy and clinical populations.

## INTRODUCTION

When Odysseus tied himself to the mast of his ship so he could hear the song of the Sirens without approaching them, he deployed a *pre-commitment* mechanism that prevented a self-control failure. When his men were unable to leave the land of the lotus eaters, Homer urges us to see them as having failed in their self-control. But what does it mean for self-control to fail? This has been a central debate in human behavior for centuries. What has fueled this debate is not a failure to understand what self-control *feels* like; the subjective experience of resisting temptation is a universal one for humans. What has made self-control so elusive is determining how to convincingly and quantitatively measure it, and therefore to understand why it often fails. Whether we are trying to lose weight, quit smoking, avoid drugs, exercise more, drink less, or simply focus on a cognitively demanding task, the question remains: If one truly desires a particular long-term outcome, why is it so difficult to choose in favor of that outcome all of the time?

Emerging theoretical accounts from psychology and economics have attempted to untangle this question using economic models of ‘cost’. Exercising self-control, these accounts propose, is *cognitively costly*. From this perspective,‘failures’ of self-control arise from a rational decision process that weighs the benefits of exercising control against its attendant costs. That is, when the cognitive costs exceed their perceived benefit, individuals should disengage from control processes. These ‘control costs’ are thought to stem from the limited cognitive resources available to support the demands of exercising control. As evidence of these costs, economic theories point to the fact that choosers often adopt *pre-commitment strategies*, presumably in an effort to reduce the need for self-control (e.g., Strotz, 1956; Thaler & Sherfin, 1981; Gul and Pesendorfer, 2001/2004). Yet, the notion of self-control as ‘costly’ remains controversial in the absence of a robust psychophysical methodology for reliably demonstrating and measuring these costs.

Historically, theoretical accounts that have attempted to explain this puzzling disconnect between what we say we want and what we actually do, have pointed to the existence of self-control without providing a platform for its reliable demonstration and quantification. The first, emerging from the psychological literature, points to self-control as a top-down regulatory process that inhibits impulsive action in the service of long-term goals or social norms. Informed by findings from classic delay-of-gratification paradigms (e.g., the ‘marshmallow task’; Mischel & Ebbesen, 1970; Mischel, Ebbesen, & Zeiss, 1972; Mischel, Shoda, and Rodriguez, 1989; but see also McGuire & Kable, 2013) and theories of ego depletion (Baumeister, Bratslavsky, Muraven, & Tice, 1998; Muraven, Tice, & Baumeister, 1998; Baumeister & Heatherton, 1996; Muraven & Baumeister, 2000; but also see Kurzban and colleagues, 2013), this account proposes that self-control relies on cognitive resources that are depleted the longer they are used. These theories suggest that the motivational or affective state of a chooser influences the availability or functional integrity of these resources. Fatigued or stressed choosers, for example, are often presumed to have more limited cognitive resources for self-control upon which to draw (Muraven, Tice, & Baumeister, 1998; Arnsten, 2009; Hockey, 1983; Holding, 1983). This body of work has dominated psychological conceptions of self-control as a form of ‘willpower’ with impulsive or suboptimal choice emerging from a failure or depletion of control resources. While this work aptly captures the subjectively difficult nature of exercising self-control, it has not provided a reliable method to quantify *how much control* is needed to successfully resist temptation.

A second account from the neoclassical economic literature examines a host of related choice problems including the failure to save money, over-consumption and procrastination (see Ariely & Wertenbroch, 2002; Bryan, Karlan & Nelson, 2010 for behavioral examples). These economic models view self-control ‘failures’ as simple preference reversals (i.e., Strotz, 1956; Mischel & Ebbesen, 1970; Ainslie, 1975; Schelling, 1984; Fishburn & Rubinstein, 1982; Bryan, Karlan & Nelson, 2010), and have generally eschewed the psychological notion of self-control as a hidden, and perhaps unnecessary, element. When individuals choose in ways that conflict with explicitly stated goals, these choosers are seen as revealing their *true preferences* through their observed choices. If this is the case however, why do individuals often choose in ways that conflict with previously stated goals, even choosing in ways that appear inconsistent or irrational (Strotz, 1955; Thaler & Sherfin, 1981; Tversky & Thaler, 1990)? Some behavioral economic work has accounted for this paradox with dual-self models that propose that choosers possess (at least) two sets of preferences that are in active competition (Thaler & Sherfin, 1981; Fudenberg & Levine; 2006), and temporal discounting models that include dynamic inconsistencies to drive changing preferences (Mazur, 1987; Ainslie, 1975; Laibson, 1997; Frederick et al., 2002; but see Kable & Glimcher, 2007/2010 for an alternate account).

While these models have provided important ways to quantify decision variables related to self-control, they do not fully explain why individuals are inconsistent in their actual choices. One widely influential resolution of this cross-disciplinary puzzle is to hypothesize that the experience of resisting immediate temptation is effortful and aversive—that is *disutile*. This inherent disutility implies that *self-control imposes a cost on choosers*, an idea formalized most recently and elegantly by Gul and Pesendorfer (2001, 2004) who proposed an axiomatic model of self-control. Gul and Pesendorfer proposed that the presence of temptation in an individual’s ‘menu’ of choices will impose a cognitive cost, rendering decisions to reject tempting options more difficult. They hypothesized that if choosers know this, they should prefer choice menus that lack tempting options and might even seek to *minimize* control costs (and maximize utility) by preemptively eliminating tempting options from their choice menu, a phenomenon referred to as ‘pre-commitment’ (Strotz, 1956; Thaler & Sherfin, 1981; see Bryan, Karlan & Nelson, 2010 for review). Examples of pre-commitment strategies that limit control costs might include a dieter who is willing to walk an extra block to avoid a local bakery or a gambler who drives an extra hour to avoid casinos. Gul and Pesendorfer thus argued that *preferences for pre-commitment reveal choosers’ subjective cost of exercising control, pointing to a novel decision variable through which these costs can be measured*.

What has limited the impact of this set of hypotheses in an empirical sense is the absence of quantitative data to support it. Is there direct quantitative empirical evidence that self-control is costly, as so many have proposed? Are those costs stationary over time? Are these costs influenced by motivational or affective state as the psychological literature has proposed? Do these costs scale with varying levels of temptation? Despite a number of real-world observations of pre-commitment (see Bryan, Karlan & Nelson, 2010 for review), we still lack an empirical psychophysical tool for answering these questions.

Here we utilized a psychophysical approach to test the hypothesis that exercising self-control is cognitively costly and that these costs can be measured using a pre-commitment mechanism. While we acknowledge that there are undoubtedly many components that feed into an overall subjective cost of self-control, our goal here was to simply measure an aggregate of these costs. To do this, we developed an economic decision-making task that measures how much participants are willing to pay to adopt a pre-commitment device that removes temptation from their choice environment. We refer to the maximum dollar value participants will pay to remove temptation as their subjective ‘cost of self-control’ and show that these costs respond rationally to incentives, scale with increasing levels of temptation and predict rates of self-control failure. We further test the hypothesis that these costs are modulated by affective state, finding that stress exposure significantly increases the cost of self-control. Finally, we test the hypothesis that self-control costs grow with the ongoing exertion of self-control but find no empirical support for this hypothesis.-These data identify a psychophysical approach for the measurement of self-control costs and may open new avenues of research into computational models of self-control to inform psychological, economic, clinical and health policy research.

## RESULTS

In our experiments, healthy, hungry dieters first provided health, taste and temptation ratings for a series of food items, allowing us to identify high- and low-tempting foods for each individual. Participants initially reported the most they would be willing-to-pay (from a $10 monetary endowment) to avoid having the high-tempting food placed immediately in front of them for a 30-minute period (**Figure 1**). With a probability of 2% this reported ‘bid’ was entered into a standard economic auction procedure (see ***Methods***; Becker, DeGroot & Marschak, 1964) that incentivizes participants to report their true subjective value—in this case the value of eliminating exposure to temptation. If they won this auction, the high-tempting food was replaced with a low-tempting food for 30 minutes. If they lost the auction, the high tempting food remained in the room with them for 30 minutes. The price participants were willing to pay provided a within-subject estimate of the cost of self-control; that is, it reflected the maximal dollar value they were willing to pay to avoid exercising control.

**Figure 1.**
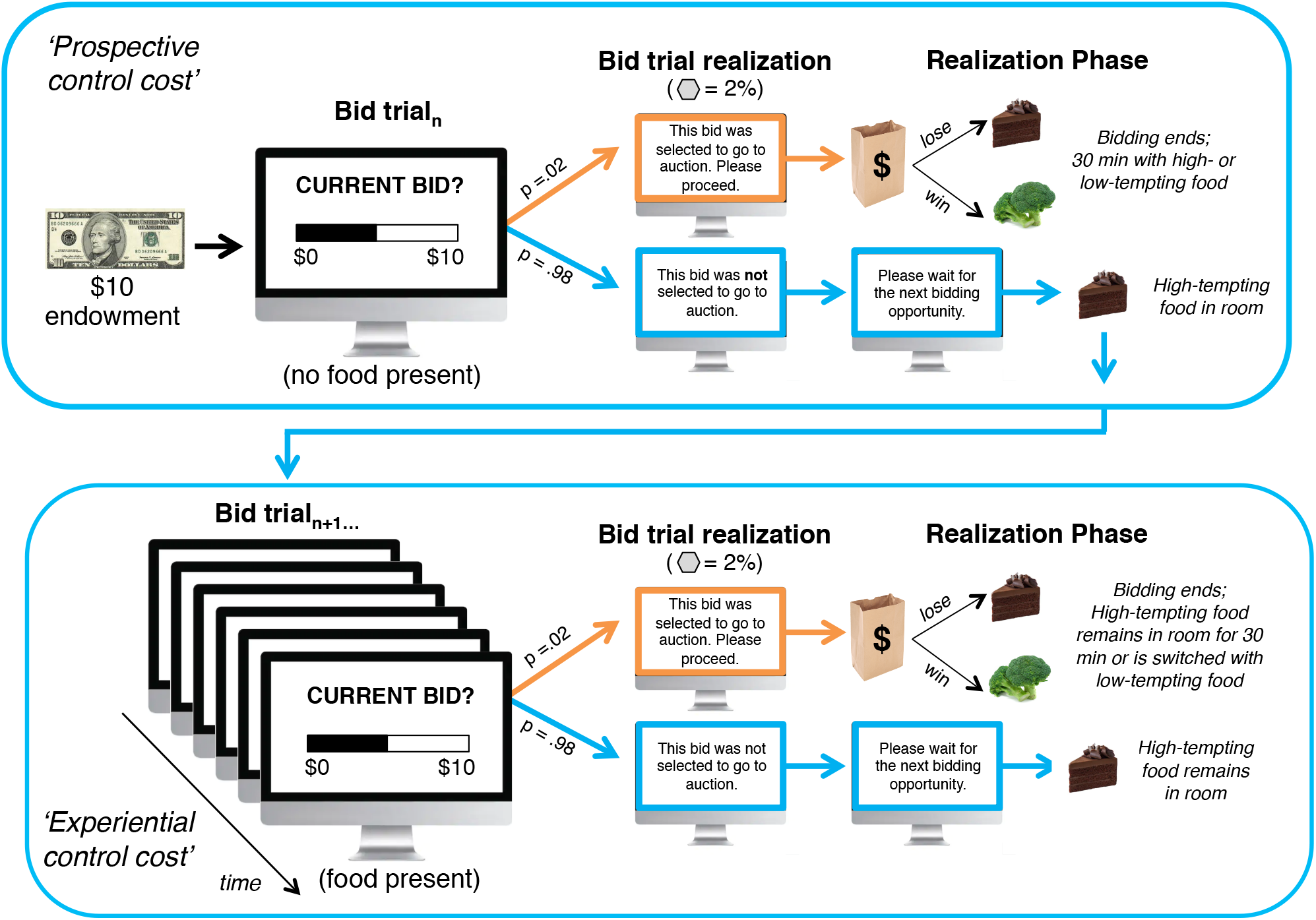
Illustration of the self-control decision task. Participants reported their willingness-to-pay to avoid a tempting food reward both before the food was present (top panel) and, periodically over a 30-minute period, with added direct exposure to the food (bottom panel).

The first bidding trial was made *without* the high tempting food in the room to capture each participants’ prospective estimate of how costly self-control exertion would be (before exposure to temptation). If the initial trial was not realized (as occurred on 98% of trials), the high-tempting food was brought into the room. Participants were then prompted to report, periodically during 30-minutes of exposure, how much they were willing-to-pay to replace the high-tempting food with the low-tempting food for the next 30 minutes. As with the initial bid, these subsequent bidding trials had a 2% chance of going to auction, which would bring the experiment to a premature close. Bidding trials were collected every few minutes (***Methods***) for the 30-minute exposure period. If the 30-minute exposure period elapsed without any bid being realized, the subjects remained in the room with the high-tempting good for a final 30 minutes. This allowed us to track how these self-control costs change over time as participants are continuously exposed to temptation and whether self-control ever failed.

The only observable behavior of interest during the final 30-minute interval of the experiment was whether or not the participant consumed the food. This *realization phase* of the experiment ensured that it was incentive-compatible, meaning participants’ choices allowed them to avoid real temptations and the negative outcomes associated with those temptations. The 2% chance of each bid being realized ensured participants knew that what they bid on the current trial was important, since it could determine whether they were required to spend the next 30 minutes alone in the experiment room with a tempting food reward, the consumption of which did not align with their stated goals.

### Study 1: Self-control imposes costs as revealed by willingness-to-pay for pre-commitment

Our primary question of interest was whether the presence of a tempting good participants want to avoid consuming does in fact impose a cost on choosers. If so, participants should be willing to pay to remove temptation and eliminate the associated control cost. In accord with predictions from these economic models, we found that participants (n=32) were willing to pay a maximum of, on average, 15% of their $10 endowment (or $1.57 ± 1.78 SD) to adopt a pre-commitment device that avoided temptation. This provides a direct scalar measurement of their subjective cost of resisting temptation (**Figure 2A)**. Participants were not only willing to pay to prospectively avoid temptation (mean of first bid trial, pre-exposure = $1.47 ± 1.59 SD), but they continued to pay throughout the task, providing a continuous measurement of the underlying costs of resisting temptation with continuing exposure to the food (mean of subsequent bid trials, post-exposure = $1.58 ± 1.82 SD).

**Figure 2.**
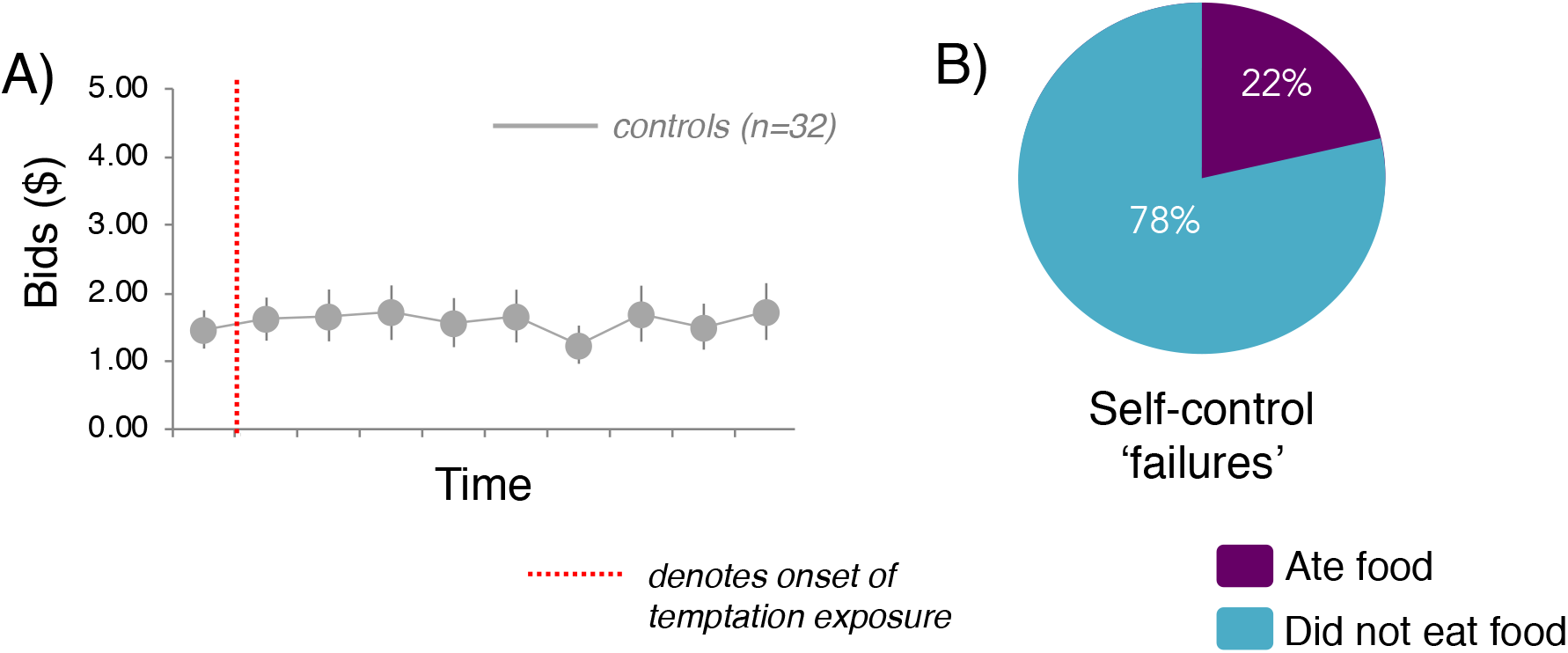
Study 1. **(A)** Bids over time for control group; **(B)** Proportion of subjects in Study 1 that consumed tempting food during the study. Error bars denote SEM.

Our initial experiment thus demonstrates that self-control imposes a subjective cost on choosers that can be measured monetarily. However, the process of deploying self-control in the presence of temptation has been proposed to change over time. Specifically, continued exposure to a tempting good is often thought to increase the difficulty of exerting self-control. We next examined the dynamics of how bids changed both before and after exposure to temptation (**Figure 2A)**. We found that average pre-versus post-exposure bids did not significantly differ (paired samples t-test: t_(31)_ = −0.533, *P* = 0.598, *d* = 0.099, two-tailed), suggesting that our dieters appear to be accurate in their prospective cost estimates. A repeated-measures ANOVA assessing post-exposure bids as a function of time indicated that on average there were no significant linear trends in these costs despite ongoing exposure to temptation (F_(4,122)_ = 0.722, *P* =0.576; Greenhouse-Geisser correction factor, *ε*=0.30; η_p_^2^ = 0.023). Thus, we found no evidence that, on average, ongoing exposure to temptation increased control costs over the interval used in our task. We note, however, that individual variability exists in our data set, such that some participants’ bids increased systematically while for others they decreased. Finally, we note that 22% of our subjects consumed the tempting food during the ensuing 30-minute exposure period (**Figure 2B**).

### Study 2: Motivational incentives modulate willingness-to-pay for pre-commitment

Next, we sought to both replicate this finding, and further test how motivational incentives to sustain goal-directed behavior might affect these costs. In this second experiment we increased the cost of self-control failure and examined whether participants were then willing to pay more for pre-commitment to reduce self-control failures. To do this, we repeated our study in a second cohort of dieters but instructed participants that they would lose a $15 bonus at the end of the study if they consumed the tempting food at any point. We hypothesized that by increasing the cost of self-control failure, the value of a pre-commitment strategy that restricts temptation should be also be higher.

Thirty-four new dieters completed our self-control measurement task with the addition of this second monetary incentive. Dieters again showed a reliable and consistent willingness-to-pay to avoid temptation (mean bid = $2.85 ± 2.70 SD). Consistent with our hypothesis, the addition of the $15 cost for eating the tempting food led to a higher willingness-to-pay for pre-commitment (**Figure 3A)**. Combining the data across experiments 1 and 2, a repeated-measures ANOVA with a within-subject factor of time (bids 1-10) and a between-subject factor of group (no incentive, incentive) revealed a main effect of group (F_(1,64)_ = 4.95, *P* = 0.03, η_p_^2^ = 0.07), but no effect of time (*P* = 0.73) or time x group interaction (*P* = 0.45). This difference at the group level suggests that participants were willing to spend more money to sustain goal-directed behavior when the costs of not adhering to this goal were higher. We note also that under these conditions none of the subjects who faced the tempting good consumed it (**Figure 3B)**.

**Figure 3.**
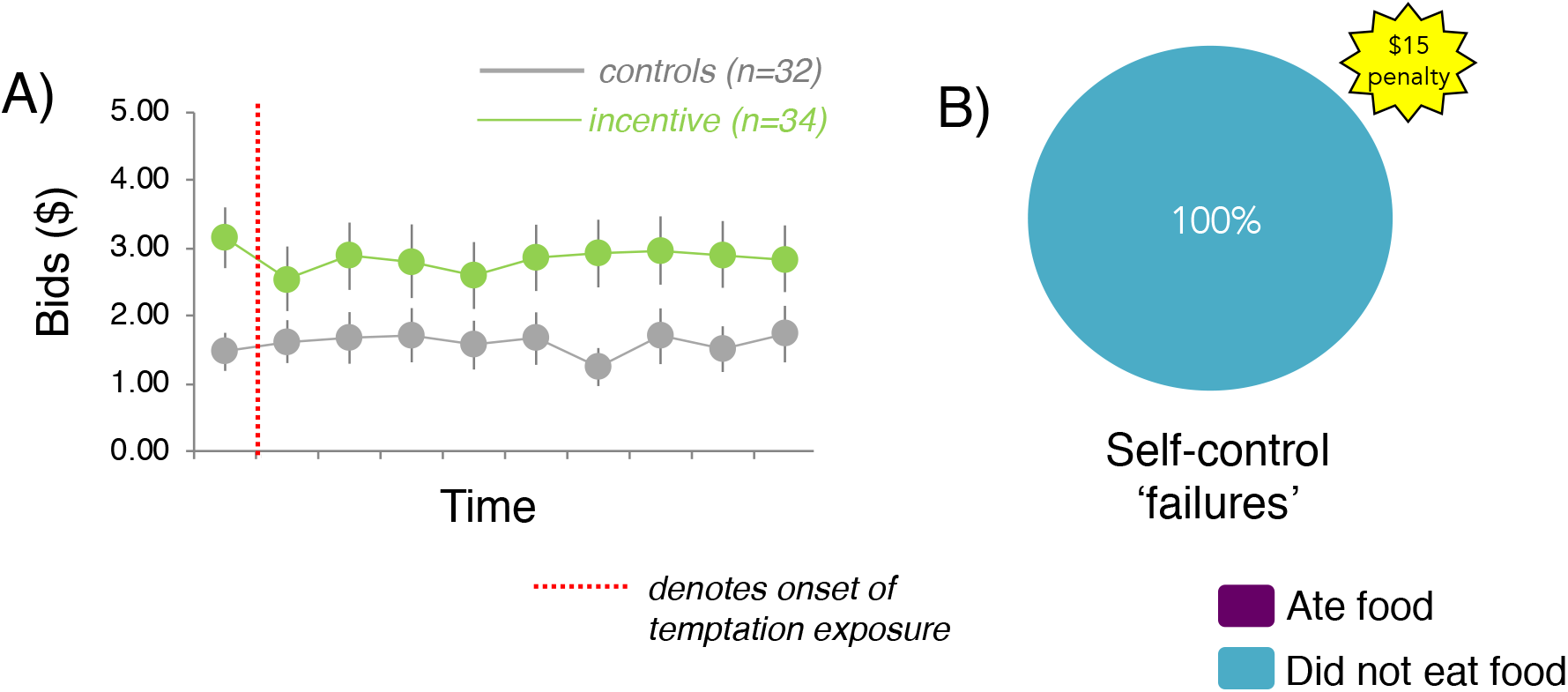
Study 2. **(A)** Bids to avoid exposure to the tempting food over time in participants for which a $15 monetary loss was imposed for consuming the food (depicted in green; Incentive group) and for those where no monetary loss was imposed (depicted in gray; Control group); **(B)** Proportion of subjects in Study 2 that consumed tempting food durin g study.

### Study 3: Acute stress increases the cost of self-control

Given the tightly coupled relationship between self-control failure and the experience of negative emotional states such as stress, we next examined how exposure to an acute stressor would influence participants’ self-control costs. Specifically, we tested the widely held hypothesis that stress makes self-control more ‘costly’. To elicit subjective and neurophysiological stress responses, we recruited a new cohort of dieters (n=31) that underwent the Cold-Pressor Task (CPT; Lovallo, 1975; Velasco, Gómez, Blanco & Rodriguez, 1997; McRae et al. 2006) prior to the self-control choice task. The CPT is widely used in laboratory settings to reliably induce mild-to-moderate levels of physiological stress and simply requires participants to submerge their forearms in ice-water continuously for 3-minutes (***Methods***). Confirming the efficacy of our stress induction procedures, participants in the CPT condition showed elevated concentrations of salivary cortisol, the primary neuroendocrine marker of Hypothalamic-Pituitary-Adrenal (HPA-) axis activity (***Figure S1A; SI Results***).

We assessed whether our stress manipulation influenced the cost of self-control both prior to temptation exposure—when participants were prospectively estimating these costs—and after the highly-tempting food was introduced. **Figure 4A** depicts aggregate trial-by-trial bidding behavior for participants in the stress condition. A repeated-measures ANOVA across all studies revealed a main effect of group (F_(2,94)_ = 4.4, *P* = 0.01; η_p_^2^ = 0.087), no effect of time (*P* = 0.45) or time X group interaction (*P* = 0.64). Follow-up t-tests confirmed that stressed participants reported a higher willingness-to-pay overall (mean bid = $3.38 ± 3.04 SD) relative to non-stressed controls (independent samples t-test: t_(61)_ =-2.88, *P*=0.005, d= 0.72, two-tailed), suggesting that our experimentally-induced state of stress was reflected in the valuation of pre-commitment to restrict temptation. These increases in the stress group were observed during prospective bids (pre-exposure, Bid1: t_(61)_ = −3.71, *P*=0.0004, d= 0.93, two-tailed) and persisted across subsequent post-exposure trials (mean of post-exposure bids: (t_(61)_ = −2.77, *P*=0.007, d= 0.67, two-tailed). Thus, exposure to acute stress appears to have more than doubled (at a between-subjects level) the average subjective cost of self-control. We note that, similar to Study 1, 23% of our subjects consumed the tempting good (**Figure 4B**).

**Figure 4.**
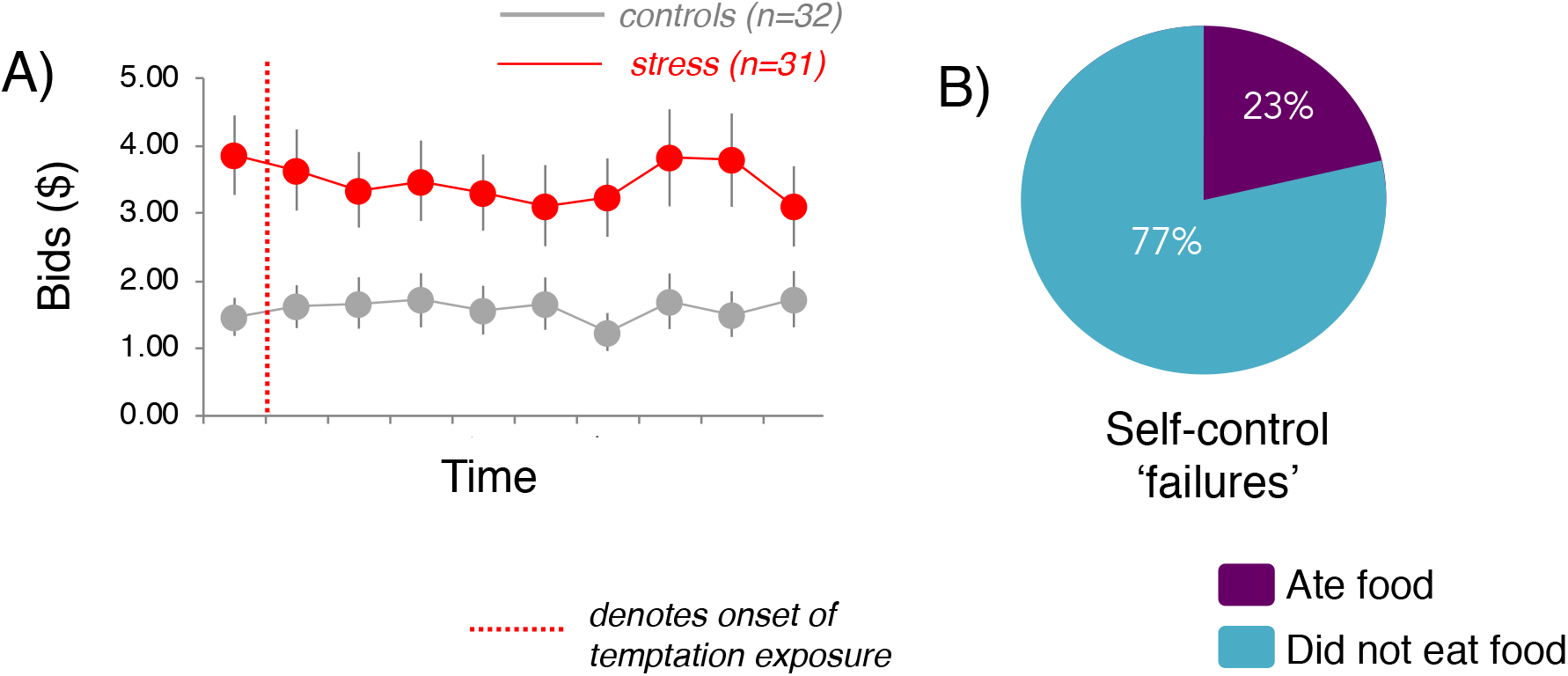
Study 3. **(A)** Bids to avoid exposure to the tempting food over time for participants that underwent a physiological stress manipulation (depicted in red; Stress group) and for non-stressed partici pants (depicted in gray; Control group); **(B)** Proportion of subjects that consumed tem pting food during sturdy.

### Study 4. The effects of stress and incentives on self-control costs

To assess whether stress exposure increased self-control costs above and beyond that which we observed when motivational incentives were introduced, we conducted an additional study on an independent cohort of dieters. Thirty-one new dieters completed the self-control decision task with the $15 penalty for eating the tempting food after undergoing the stress manipulation. We once again observed a reliable willingness-to-pay to avoid temptation (mean bid = $2.74 ± 2.20 SD) that was elevated relative to the Study 1 controls (see below and ***Figure S2, SI Results***; *t*_(61)_ = −2.34, *P* = 0.023, two-tailed). To assess if stress exposure changed bids relative to non-stressed participants who experienced the same incentive structure, we conducted a time X group RM-ANOVA. However, this analysis revealed no main effect of time (*P* = 0.55) or group (*P* = 0.86), nor time x group interaction (*P* = 0.98), thus revealing no significant differences in bidding behavior (***Figure 5***). We reasoned that a failure to observe group differences as a function of stress exposure could be due to the fact that the stress manipulation in this particular cohort did not effectively elicit cortisol responses. Indeed, an examination of cortisol concentrations revealed that cortisol levels in this cohort did not differ between groups at any time point (***Figure S1B, SI Results***). Thus, it appears that while motivational incentives led to an increase in bids relative to control participants (replicating our effects from Study 2) we did not observe differences between the two incentive groups as a function of stress exposure, perhaps due to the failure of the stress procedure.

**Figure 5.**
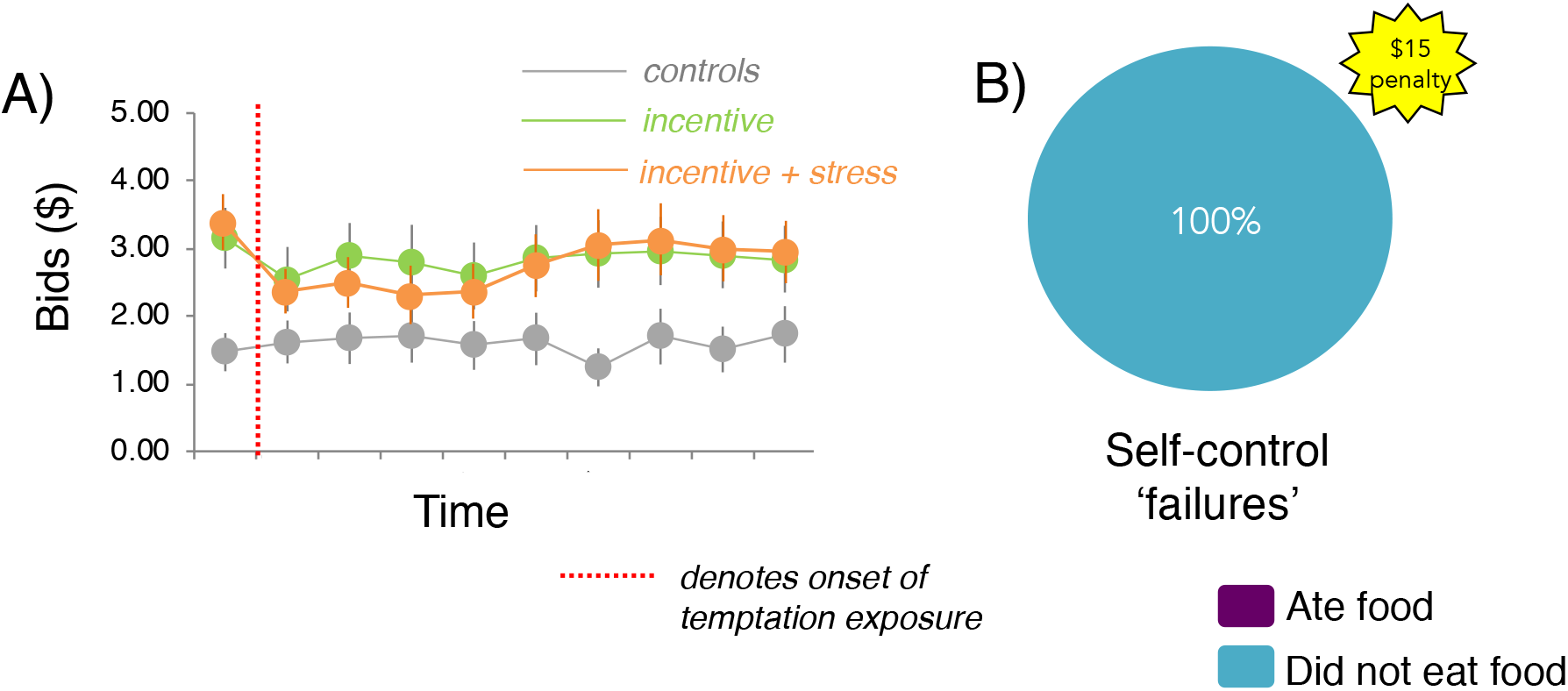
Study 4. **(A)** Bids to avoid exposure to the tempting food over time in stressed partici**p**ants for which a $15 monetary loss was imposed for consuming the food (depicted in orange; Incentive+Stress group) and for those with the same penalty imposed but no stress insluction (depicted in green; Incentive group; Study 2). The control group from Study 1 (no incentive or stress) is depicted in grey for reference. **(B)** Proportion of subjects from Study 4 that consumed the temptinţg food during the study.

### Secondary Analysis 1: Self-control ‘failures’ were associated with a higher willingness-to-pay to avoid control

If we assume that subjects do face costs for exercising self-control, then we should expect to see subjects experience self-control failures on occasion. Further, we might expect to find that subjects willing to pay more for pre-commitment experience higher self-control costs and thus might be expected to fail in their self-control more often than subjects who report lower monetary costs for self-control. To test these hypotheses, we examined each subject’s behavior after the bidding phase of the experiment was complete—during the final 30-minute phase of the experiment (***Methods***). During this phase, if no bids had been realized (which was usually the case given the low probability any bid trial was realized), participants were required to remain in the room for the final 30 minutes of the experiment with the tempting food. No further bids were collected during this phase. During this period, we simply observed whether or not each participant consumed the tempting food (a self-control “failure”). The proportion of participants that consumed the tempting food are presented alongside bidding behavior (**Figures 2B–5B**). We focused our comparison on the study cohorts that shared similar penalty structure in order to control for the increased cost of self-control failure. In Study 1 and 3, where no monetary penalty was imposed for eating the tempting food, 22% and 23% of participants consumed the food, respectively (**Figure 2B and 4B**). Given that consumption rates did not differ between these two groups, we collapsed across these studies to examine how bidding behavior differed in dieters that ate the food vs. those that did not. Those participants who ate the food were willing-to-pay significantly more to avoid temptation relative to participants who did not eat the food (independent samples t-test: t_(61)_ = 2.81, P=0.006; two-tailed; **Figure 6**). In Study 2 and 4, where we imposed a monetary penalty for self-control failure, no participants consumed the food (**Figure 3B and 5B**), thus a comparable analysis to that of above was not possible. We note that the above differences between ‘eaters’ and ‘non-eaters’ remain significant even when including all ‘non-eater’ participants from Study 3 (t_(95)_ = 2.41, P=0.02) and, additionally, Study 4 (t_(126)_ = 2.43, P=0.02).

**Figure 6.**
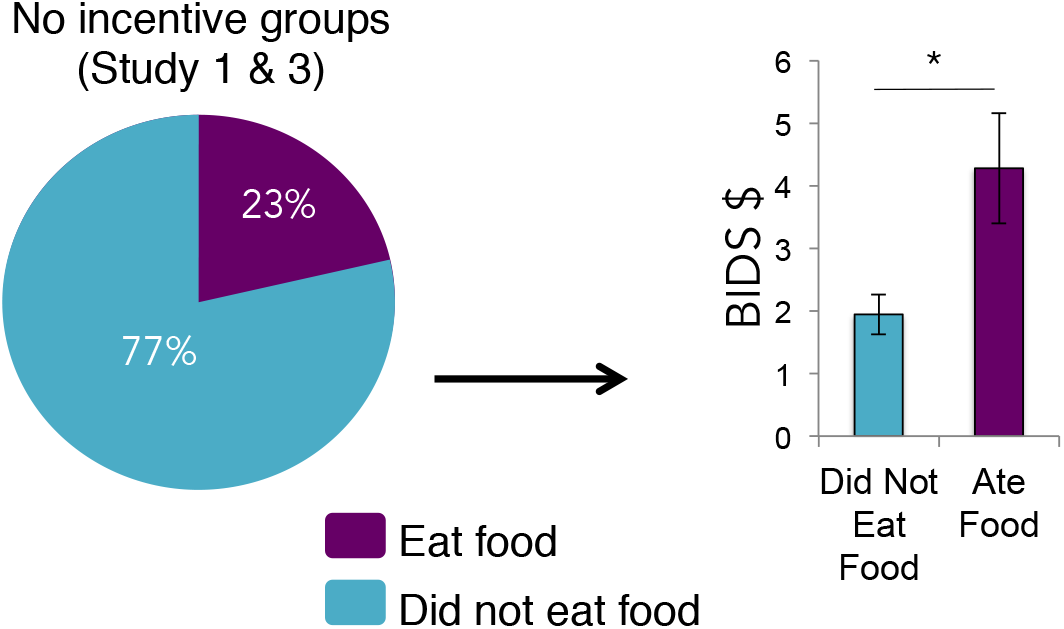
Mean bid for subjects who demonstrated self-control ‘failures’ (23%, depicted in purple) and those who did not (77%; depicted in blue) collapsed across Study 1 (control group) and 3 (stress group). Those participants who consumed the tempting food during the study revealed a higher willingness-to-pay to avoid control. No participants from Study 2 or 4 ($15 penalty groups) consumed the tempting food. Error bars denote SEM.

### Secondary Analysis 2: Individual difference measures associated with control costs

We next examined how individual differences across our entire sample related to self-control costs. Given that control costs were higher in stressed participants (Study 3), we first sought to characterize how subjectively perceived stress related to willingness-to-pay to avoid self-control. To do this, we correlated mean bids and self-reported stress levels directly before the choice task across participants from all four studies (n=128). Perceived stress was positively correlated with average bidding behavior (Spearman’s rho: *r* = 0.22, *P* = .012; **Figure 7A**), suggesting that, across all participants, subjective stress state was related to individuals’ willingness-to-pay to avoid temptation.

**Figure 7.**
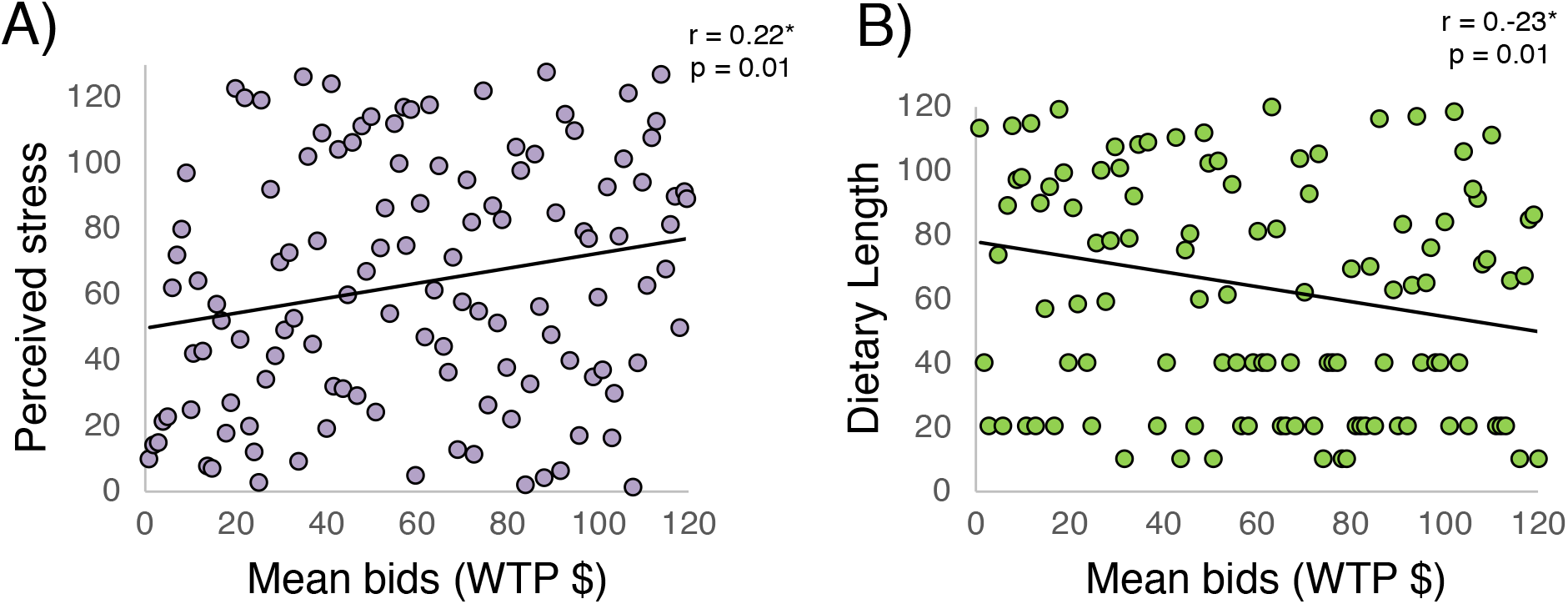
Individual Difference Correlations. **(A)** Perceived stress was positively correlated with average bidding behavior across participants; **(B)** Length of diet was n egatively correlated with average bidding behavior across participants. *\*denotes Spearman ranked correlation*

Given our evidence that self-control is explicitly costly to choosers, we reasoned that more experience (or success) avoiding temptation might relate to an individual’s self-control costs. To explore this question, we conducted a correlation analysis between participants’ length of diet and their average willingness-to-pay to avoid exercising self-control. This analysis revealed a significant negative correlation between mean bids and diet length (Spearman’s rho: *r* = −0.23, *P*=.01; **Figure 7B**). Thus, participants on a diet for a longer length of time tended to pay less to avoid temptation. We note that this does not reveal whether only those with idiosyncratically lower self-control costs succeed at remaining on a diet, whether self-control costs decline as one’s diet progresses, or both. However, our method applied longitudinally could be used to answer such questions in future work.

### Study 5: Willingness-to-pay to avoid control scales with temptation level

If participants’ willingness-to-pay to avoid temptation does in fact reflect the cost of self-control, then we would expect these costs to scale with varying levels of temptation (i.e., when facing a more highly tempting good, a subject should have to exert more self-control than when facing a less tempting good). To test this, we conducted a final study in an independent cohort of healthy, hungry dieters. In this study, participants again rated a series of snack foods on how healthy, tasty and tempting they were, which allowed us to identify a low, medium and high tempting food for each individual. On each trial, participants reported their willingness-to-pay (from $0-$10, from a $10 endowment) to avoid each of the three food items for varying amounts of time (1-60 minutes; ***Methods***). Unlike Study 1 −4, *all* bids were reported prospectively (without any food present) and one trial was randomly selected at the end of the session to be realized.

**Figure 8** depicts average bids for each time point for each level of temptation. A temptation level (low, medium, high) X time (1-60 minutes) RM-ANOVA revealed a significant main effect of temptation level (*F*_(2,30)_ = 33.06, *P* < .0001, *η_p_^2^* = 0.69) and time with food (*F*_(9,135)_ = 29.12, *P* < .0001, *η_p_^2^* = 0.67), as well as a temptation X time interaction (*F*_(18,270)_ = 5.75, *P* < .0001, *η*_p_^2^ = 0.27). Bids differed significantly for foods with low (mean bid = $1.16 ± 0.36 SD), medium (mean bid = $2.99 ± 1.02 SD), and high (mean bid = $4.96 ± 1.30 SD) temptation levels. Further, bids scaled with each temptation level across increasing amounts of time (**Figure 8**). Planned contrasts demonstrated a significant linear increase in bids with higher temptation level (*F*_(1, 15)_ = 43.95, *P* < .0001) and, separately, with increased time with food (*F*_(1, 15)_ = 44.81, *P* < .0001).

**Figure 8.**
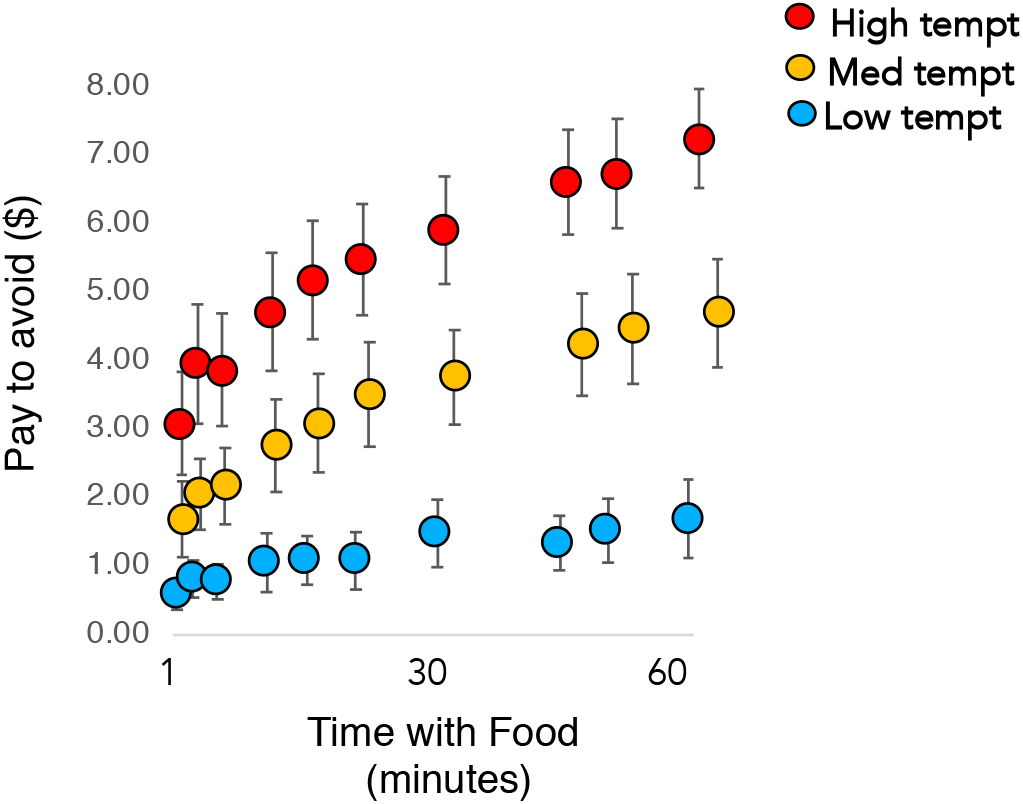
Willingness-to-pay to avoid foods that varied in temptation level and amount of time required to spend with food. Bids scaled with increasing time with and temptation of the food. Error bars depict SEM.

## DISCUSSION

A universal paradox in human behavior is our tendency to set difficult long-term goals but then to make choices that appear to contradict or undermine those goals. Using an economic decision-making paradigm, coupled with a sample of dieters avoiding tempting food rewards as a model of self-control more broadly, we found that people will pay to avoid temptation, quantitatively revealing their subjective cost of control under a variety of circumstances. We found that these costs are modulated by incentives (shows cost sensitivity; Study 2/4) and acute stress exposure (Study 3), consistent with the notion that motivational and affective state modulate one’s willingness to exercise self-control. In a final study designed to test how self-control costs scale with increasing levels of temptation (Study 5), we showed that longer exposure to a tempting good imposes higher self-control costs (self-control costs obey monotonicity) and more tempting goods impose higher control costs at the within-subject level than do lower tempting goods (self-control costs order with temptation).

Decades of psychological research have revealed that the act of engaging in self-control is subjectively effortful and aversive. Emerging work in the cognitive control literature has proposed that this experience stems from the cognitive costs imposed by deploying control, an account consistent with more recent economic theories of self-control (Gul & Pesendorfer, 2001/2004) that propose preferences for pre-commitment reveal an inherent psychological cost to resisting temptation. These converging lines of work provide a clear and testable hypothesis: Self-control failures may be conceptualized as a rational decision that emerges when the costs of exercising control exceed the relative perceived benefits (Kurban et al., 2013; Berkman et al., 2017; Kool & Botnivick, 2018). However, without a psychophysical method to precisely quantify these costs, researchers often must *infer* whether and how much self-control a chooser requires to make goal-consistent decisions or to delay gratification. Our findings unite psychological and economic theories of self-control and provide empirical evidence that self-control is explicitly costly to humans and that these costs can be quantified by measuring the value of pre-commitment to restrict temptation.

Gul and Pesendorfer’s theory is consistent with a growing body of psychological and neuroscience research that suggests people view cognitive demand as intrinsically costly and tend to avoid utilizing cognitive resources if possible (Kurzban et al. 2013; Westbrook & Braver, 2015; Shenhav et al. 2013/2017; Kool & Botnivik, 2018). These costs are thought to stem from limitations in the cognitive resources available to support the demands of control, suggesting self-control arises from evaluations of how valuable expenditures of control are perceived to be relative to how costly. This is a notable departure from classic psychological self-control models that view cognitive resources as depletable (i.e., both limited and diminished with use), arguing instead that such resources are finite and reallocated dynamically depending on the perceived costs and benefits (Kurzban et al., 2013; McGuire & Kable, 2015). Why some classes of cognitive control may be aversive or costly remains unclear. However, the approach presented here offers a metric for how aversive or costly individuals’ find the exertion of self-control to be on a moment-to-moment basis.

Our task employed two important features to probe the nature of self-control. First, our task measured momentary willingness to pay to avoid control prospectively (prior to food exposure) and again after participants encountered the tempting food. One possible explanation for why self-control appears to fail so often is that individuals may poorly estimate how costly self-control will be. Our data tends to lean against this conclusion, at least in this particular choice setting. We observed no significant difference in measured control costs before versus after food exposure. This suggests that our participants had an accurate prospective awareness of the self-control costs they would later face. A second important feature of our task was that—unlike existing self-control decision paradigms—our bidding measurements were collected continuously over time, allowing us to track how these costs change with continued exposure to temptation. Some existing work (Baumeister, Bratslavsky, Muraven, & Tice, 1998; Muraven & Baumeister, 2000) suggests that that self-control becomes more difficult as it is continuously exerted. If this were true for our participants, we would expect their self-control costs to increase as the experiment progressed. Interestingly, we instead observed that participants’ bids to avoid control were quite stable over time. The fact that participants’ control costs did not increase over time, however, is consistent with a growing body of work suggesting that performance reductions are a not mandatory feature of engaging in control (Kurzban et al., 2013; Shenhav et al. 2013/2017; Kool & Botnivik, 2018). Our findings are consistent with value-based frameworks that argue individuals need not necessarily experience decrements in control performance as long as the perceived benefit of deploying control continues to outweigh the cost. We note, however, that our data on this point is relevant only to the intervals of time tested here (under an hour in total duration).

The stability in our participants’ willingness to pay to avoid exerting control may also reflect the *lack of temporal uncertainty* inherent in our task. In a recent line of work, McGuire and Kable (2013/2015) demonstrated that behavior in a range of delay-of-gratification tasks (including the ‘marshmallow task’, Mischel & Ebbesen, 1970) might be explained by the underlying predictions participants have regarding when a delayed outcome will arrive. They suggested that one major reason individuals appear to ‘succumb’ to temptation is that under temporal uncertainty, individuals may rationally conclude that the delayed outcome may no longer be worth waiting for. In our study participants were fully informed regarding the temporal structure of the task and were at all times aware that self-control would be engaged only for a finite period of time. This feature of our task may explain the stability observed in participants reported bids over time. An open question for future research is whether imposing temporal uncertainty, or requiring self-control for longer periods of time (Blain et al., 2016) than we used in our task, might lead to an increase in ongoing self-control costs even at this limited time interval.

Self-control research across disciplines suggests that we should be able to induce changes in these costs with changes in motivational state. Consistent with the notion that decisions to engage in self-control arise from a dynamic cost-benefit evaluation (Berkman et al., 2016), we found that willingness to pay for pre-commitment increased as the cost of failing to adhere to one’s diet increased. When faced with losing $15 in addition to failing to adhere to their diet, participants were willing to pay to restrict access to temptation, thus pre-commitment became more valuable (Study 2 and 4). These findings are consistent with work showing that motivational incentives can increase willingness to engage in self-control strategies (Hagger et al, 2010; Krebs et al., 2010) and demonstrate overall cost-sensitivity in the self-control mechanism.

We also observed an increase in the cost of self-control when participants were successfully stressed using an acute stress induction (Study 3). Stress exposure has long been thought to compromise self-control (Hockney, 1983; Holding, 1983), and this intuition has been borne out in a large body of empirical work— from the cognitive neuroscience literature that shows stress diminishes cognitive capacity and flexibility (Schoofs et al 2009; Plessow et al 2009) and selectively reduces goal-directed control of decisions (Schwabe & Wolf 2009; Otto, Raio, et al., 2013) to the clinical literature where stressors remain a primary risk factor for the emergence and relapse of addiction-related disorders (Sinha, 2011). Our findings provide a direct test of whether stress compromises self-control by increasing its associated cognitive cost. This relationship was also observed beyond participants assigned stress condition, as higher self-control costs were positively associated with perceived stress scores. We note that in Study 4, where both stress and incentives were imposed before the self-control task, we did not observe an added increase in bids above and beyond that of the Study 2 incentive group. However, this may be due to the fact that we did not see evidence of a cortisol elicitation from the stress manipulation. Future work may seek to use alternate stress induction techniques (e.g., social or cognitive stressors) to test if other types of stressors lead to additive (or interactive) effects with motivational incentives on self-control costs. Finally, one can imagine decision environments whereby stress can impair pre-commitment decisions given their reliance on prospective thinking. Our task utilized both prospective and experiential exposure to temptation but it would be interesting to examine how stress affects pre-commitment decisions that rely fully on future prospection or memory retrieval.

Finally, in Study 5, confirming that our bids did in fact reflect the cost of self-control and not random baseline bidding behavior, we found that the average magnitude of bids scaled with increasing levels of temptation. By testing bidding behavior across varying degrees of temptation level and a broader range of times with the food, we were able to confirm that willingness to pay for pre-commitment tracks with the increased cost of resisting temptation, demonstrating that the cost of self-control appears to increase monotonically with duration and that more highly tempting goods induce higher self-control costs. Overall, our findings suggest that measuring the control costs can reveal unique information about the subjective experience of resisting temptation that cannot be attained using existing measures of self-control.

Given its importance as a strategy to help individuals achieve their long-term goals by reducing self-control costs, a number of recent studies have begun to measure preferences for pre-commitment in the laboratory (Crockett et al., 2013; Schwartz et al., 2014; Soutschek et al., 2017/2020; Studer et al., 2019). For example, Crockett et al. (2013) provided the first empirical demonstration that pre-commitment facilitates choices for larger, later rewards as opposed to smaller, sooner ones when explicitly offered as a choice strategy in an intertemporal choice task (e.g. viewing erotic images that varied in reward magnitude and delay). This study further demonstrated that the primary neural circuits underlying pre-commitment (e.g., lateral frontopolar cortex; also see Soutschek et al., 2017) are distinct from that of standard self-control (“willpower”) tasks that rely on the effortful inhibition of impulses (e.g., dorsolateral PFC, inferior frontal gyrus), consistent with the notion these self-control strategies rely on different neurocognitive processes. These data, coupled with more recent work showing that higher impulsivity and meta-cognitive awareness leads to stronger preferences for pre-commitment (Soutschek et al., 2020), are consistent with the view that pre-commitment decisions engage future planning and prospection and are driven by an awareness of subjective self-control costs.

Our results add to this growing literature by demonstrating that individuals not only show preferences for pre-commitment, but they are willing to pay to adopt such strategies, effectively revealing their subjective cost of self-control over time and with continued exposure to temptation. Our task extends extant studies of pre-commitment outside of classic intertemporal and effort-based choice paradigms by directly quantifying the cost of resisting temptation under different motivational/affective states and levels of temptation. One relevant question for future work is how we can further dissociate the cost-benefit processes that drive pre-commitment decisions. For example, future work might address to what extent individuals pay to remove temptation in order to avoid a self-control failure, versus to diminish the disutility of resisting temptation irrespective of predicted failures. One recent study, for example, found that pre-commitment increases motivation to engage in effortful action that leads to larger rewards. Specifically, in the context of an effort- and delay-based choice task, pre-commitment choices were found to be driven by a desire to reduce opportunity costs and secure adequate motivation to endure the longer delay or higher effort required to attain larger rewards, rather than to avoid a failure of willpower per se (Studer et al., 2019). Future work may attempt to further dissociate the motivational processes that underlie these decisions and how changes in temptation intensity or temporal uncertainty may alter them.

A number of limitations should be noted for future work. First, we acknowledge that there are likely many different components to what makes the exertion of self-control cognitively costly. For example, there may be cognitive costs to resisting temptation and also personal and health costs associated with self-control failures. While our task does not currently dissociate among the components that feed into self-control costs more generally, this is an open area for future research. Second, unlike some studies in the human stress literature we included both men and women in each of our samples. However, we did not measure menstrual phase, oral contraceptive use or cycle-dependent sex hormones, which have been shown to impact stress responses in women (Kirschbaum et al., 1999; Hellhammer et al., 2009). Future work measuring such factors may reveal interesting patterns in control costs that we were unable to detect here. Finally, despite every effort to eliminate them from our procedure (*SI Materials*), we cannot definitively rule out the possibility that demand characteristics may have contributed to these effects in some way.

In summary, we report a novel task for measuring the subjective cost of self-control. Our findings are consistent with emerging work across disciplines suggesting that self-control and its failures can be seen as fundamentally rational responses to a complex world in which individuals’ trade-off the cognitive cost of resisting immediate temptation against the benefits of achieving future goals.

## METHODS

### Participants

138 healthy young adult participants that indicated they were on a diet to maintain or lose weight participated in the study (see ***SI Materials*** for full screening criteria). Participants were recruited using flyers posted on and around the NYU campus, as well as electronic advertisements on New York University’s Department of Psychology website. Participants were excluded prior to participation for the following reasons: (1) pregnancy; (2) high-blood pressure or a heart condition; (3) history or medication for neurological or psychiatric disorders; (4) diabetes, metabolic disorders, food allergies or history of eating disorders; and (5) use of corticosteroids or beta-blockers. All participants provided written informed consent in accordance with experimental procedures approved by the New York University Committee on Activities Involving Human Subjects. All research and experimental procedures were performed in accordance with approved IRB guidelines and regulations.

Subjects were paid $15 per hour plus a $10 bidding endowment. Six participants from the stress groups were unable to complete the CPT task and were thus excluded. Two additional participants were removed prior to data analysis because they revealed that they were on special diets to sustain (and ideally increase rather than decrease) weight and two others were excluded for being on medication (revealed after the experiment ended). Our final analysis included a total of 128 healthy participants (84 women) with a mean age of 24.37 (SD = 7.07; range = 18-55).

### General Procedure (Study 1-4)

Hungry, healthy dieters were asked to abstain from eating 3-4 hours before participating in the study. Upon arrival at the laboratory, participants provided informed consent and were escorted to the experiment room for a 10-minute acclimation period, after which they rated their current hunger level (from 1-10; **Fig. S3 and S4**, ***SI Results***), completed the food rating and ranking scales (**Fig. S5**, ***SI Results***) and provided basic information about their current diets (**Fig. S6**, ***SI Materials***). After baseline cortisol was collected, participants received their $10 (cash) endowment. They then received instructions regarding the self-control decision task and were explicitly informed about which high- and low-tempting foods they would be making choices about during the study. (Participants in Studies 2 & 4 were further instructed that they would lose a $15 bonus payment provided at the end of the study if they consumed the tempting food at any point.) All participants then completed either the CPT or control task and were given a 10-minute break in the experiment room before an additional cortisol sample was collected. This break was implemented to ensure that cortisol levels induced by the CPT would begin to peak in coordination with the choice task. Participants then completed the self-control decision task (see ***Decision-Making Task***), during which they indicated the maximum amount that they would be willing to pay to remove the high-tempting food from the room and replace it with the low-tempting food. After the 30-minute bidding phase of the task was complete, the final cortisol sample was collected and the final phase of the experiment began, during which participants were required to remain in the experiment room with the high- or low-tempting food for the final 30 minutes of the study (see ***Bid Realization***). Once this 30-minute final phase was over participants were paid for their time and left the laboratory.

### Stress Induction Technique

All participants in the stress group (Studies 3 & 4) completed the CPT, for which participants submerged their right forearms in ice-cold water (0 °C to 4 °C) for 3 min continuously. All participants in the control group (Studies 1 & 2) followed the exact same procedure using room-temperature water (30 °C to 35 °C). The CPT is widely used in laboratory settings to model the effects of mild to moderate stress and reliably generates both autonomic nervous system and HPA-axis activation, as measured by increased physiological arousal, neuroendocrine responses, and subjective stress ratings (Lovallo, 1975; Velasco, Gómez, Blanco & Rodriguez, 1997; McRae et al. 2006).

### Neuroendocrine Assessment

Saliva samples were collected throughout the study to assess cortisol concentrations, which serve as neuroendocrine markers of stress response. Participants were run between 12 and 5pm to control for diurnal rhythms of stress hormone levels. Saliva samples were collected using a high-quality synthetic polymer-based salivette placed under participants’ tongues for two minutes. Participants were initially seated in the experiment room for a 10-minutes acclimation period, during which they drank 4 oz of water to clear any residual saliva. Samples were collected at baseline (sample 1), 10 minutes after the CPT/control task administration (sample 2), and directly before the choice realization began (~30 minutes after the CPT/control task administration; sample 3). Samples were immediately stored in a sterile tube and preserved in a freezer set to −80 °C. Samples were analyzed by the Psychobiological Research Laboratory of the University of Trier, Germany, using a time-resolved immunoassay with fluorometric detection (DELFIA, cf. Dressendörfer et al., 1992). Duplicate assays were conducted for each sample, and the average of the two values was used in our analyses. Any samples that contained insufficient saliva could not be analyzed and were excluded from our analyses. Cortisol data was log-transformed to account for the skewed nature of cortisol distributions (**Fig. S1A and S1B**, ***SI Results***).

### Food Item Scales & Selection

In order to select a high- and low-tempting food item for each individual, participants completed a series of food rating scales prior to the study (**Fig. S5**, ***SI Results***). Participants separately rated 20 food items (**Fig. S7**, ***SI Materials***) on how tasty, healthy and tempting these items were from 1 (not at all) to 10 (very much so). Participants then ranked these 20 food items from best (#1) to worst (#20) for their current diet. Low-tempting foods were chosen by a computer algorithm that identified foods that fell in the lowest 20% of taste and temptation ratings, the highest 20% of health ratings and that was ranked in the upper 10% of foods best for the participants’ current diet. Conversely, high-tempting foods were identified as those that fell in the upper 20% of taste and temptation ratings, the lowest 20% of health ratings and that was ranked in the lowest 10% of foods worst for the participants’ current diet.

### Decision-making Task (Figure 1)

To directly examine individuals’ subjective cost of self-control, we designed an incentive-compatible decision-making task that measured the monetary costs that participants were willing to incur to avoid temptation on a moment-to-moment basis. There were two phases of the task: (1) a bidding phase, during which participants indicated the maximum they would be willing-to-pay from a $10 endowment to remove the high-tempting food from the room and replace it with the low-tempting food; and (2) a final realization phase, during which participants sat in the experiment room with the food for the final 30 minutes of the experiment. Participants in Studies 2 & 4 were further instructed that they would lose a $15 bonus payment at the end of the study if they consumed the tempting food at any point.

On each trial, participants viewed a computer screen that prompted them to enter the maximum amount that they were currently willing-to-pay to remove the high-tempting food from the room and replace it with the low-tempting food. Participants registered their bids with an open response-window by using the mouse to control a sliding bar that ranged from $0 to $10 (in $0.01 increments) and clicking the mouse on their selected bid. Bid trials were presented approximately every 3 minutes (Studies 2 & 4) or 2 minutes (in an effort to acquire more precise temporal measurements of bids) for a total of 10 bids for Study 2 & 4 and 15 bids for Studies 1 & 3. Choices were presented using PsychToolBox.

After each bid was received there was a 2% chance that this bid would be immediately employed in a procedure that would lead to the final 30-minute phase in which the high-tempting or low-tempting food would be in the room with the subject. This incentivized participants to bid their true value for removing the high-tempting food since the bidding phase of the task could end on any trial and only the current bid would be used to determine whether the food was removed for the 30-minute final phase. The 2% hazard rate also ensured that the majority of bidding trials would not be realized, allowing us to track the dynamics of how self-control costs change over time with greater exposure to temptation. Finally, this feature of the task allowed us to eliminate any effect of temporal discounting on the sequential bids. To realize bids, we used a standard Becker-DeGroot-Marschak (BDM) auction procedure widely used to reveal maximum willingness-to-pay (see ***Bid Realization Procedure*** below).

For the initial bidding trial, no food was present in the room. If this initial trial was not realized, the food was brought in the room and remained there until a bid was implemented or the bidding phase concluded. (If the trial was realized the BDM procedure determined whether the food was brought into the room for 30 minutes. Immediately after each bidding trial, participants were notified as to whether that particular bid would be implemented. If no bid was realized by the end of the 30-minute bidding phase, the task transitioned automatically to the final 30-minute realization phase.

### Self-Control Failures

Any quantity of food consumed during the task was considered a self-control failure. In all cases of eating, participants consumed the entire item with the exception of one participant who consumed half of the snack food (potato chips) presented. Any participants that consumed the food did so during the final 30-minute realization phase. To reduce observer effects, participants were alone in the experiment room for the duration of the study so we were not present to measure the precise time at which they might have eaten the tempting food. Whether or not they consumed the food was revealed after participants left the laboratory.

### Bid Realization Procedure

To determine whether the participant won or lost the opportunity to replace the high-tempting food with a low-tempting food, a standard economic auction procedure was implemented (DeGroot-Marschak; BDM). Participants selected a selling chip from a bag (chips ranged from $0 to $10 in $0.01 increments) and this selected chip represented the *winning sell price*. This randomly selected sell price was then compared against the participant’s current bid. If the bid price was greater or equal to that of the sell price, then the participant won the auction and they paid the *sell price* (not the bid price) from their endowment to have the high-tempting food removed. If the bid was lower than the sell price the participant lost the auction. In this case, the high-tempting food remained in the room for the remainder of the experiment and the participant would keep the entire endowment. This procedure incentivizes participants to report their true maximum willingness-to-pay.

### Study 5

An additional 20 participants were recruited following the same recruitment, screening, informed consent and payment procedures as Study 1-4 (***Participants***). Two participants were excluded for using medications on our exclusion criteria list. Due to a computer software error, data from two other participants was not recorded. Participants were asked to refrain from eating prior to coming into the lab and began by rating the same series of 20 food items (see ***Figure S7*** for choice set) on how tasty, healthy and tempting these items were from 1 (not at all) to 10 (very much so). These ratings allowed us to select a low, medium and high tempting food for each participant. On each trial, participants viewed an image of a snack food that varied on temptation level (low, moderate, high) and amount of time for which participants would potentially have to spend with the food (1, 3, 5, 10, 15, 20, 25, 30, 45, 60 minutes). Participants viewed the food for 4 seconds and the entered their how much they were willing to pay (from $0-$10) from a $10 endowment to avoid that food, given the temptation level, quantity and time amount. After the 90 trials were completed, one trial was randomly selected and the same BDM auction procedure was used to identify whether the participants won or lost the auction based on their bid for that given trial (***Bid Realization Procedure***). All participants remained in the experimental room for 1 hour after the bidding task was complete in order to control for the cost of time (i.e., to ensure bids did not reflect an aversion to waiting for the allocated amount of time relative to the cost of control). If participants lost the auction, the food was present in the room for the amount of time stated on the selected trial (e.g., if the trial depicted 15 minutes with the food, then the participant spent 15 minutes of the full hour with the food). If they won the auction then the food was not present during this amount of time.

### Data Analysis

All statistical analysis for behavioral and cortisol data was carried out using SPSS (version 20.0, 2011; IBM Corp., Armonk, NY). Due to the skewed nature of cortisol concentration distributions documented in the literature (Miller et all, 2013), cortisol values were log-transformed in order to better approximate a Gaussian (normal) distribution. Data were tested for equal variances using Mauchly’s sphericity tests and Greenhouse-Geisser corrections were performed to address any violations of sphericity. Analysis of variance (ANOVA) with repeated measures was used to analyze all choice (i.e., bidding) and cortisol data. Post hoc comparisons were conducted using Student *t*-tests when appropriate. All tests were two-tailed and considered statistically significant when p < .05.

## Data and Code Availability

The data sets generated during and/or analyzed during the current study are available from the corresponding author on reasonable request.

## Acknowledgments

We thank all members of PWG‘s laboratory, especially Anna Konova, Alexandra Mellis and Christopher K. Steverson, for helpful feedback, comments and discussion. This work was supported by NIH Grant F32MH110135 and a NARSAD Young Investigator Grant from the Brain & Behavior Research Foundation to CMR and NIH Grant R01DA038063 to PG.

## Notes

### Competing Interest Statement

The authors have declared no competing interest.

### Summary of Updates

Clarifications throughout; additional figures and updated Supplemental files.

